# Identifying and removing haplotypic duplication in primary genome assemblies

**DOI:** 10.1101/729962

**Authors:** Dengfeng Guan, Shane A. McCarthy, Jonathan Wood, Kerstin Howe, Yadong Wang, Richard Durbin

## Abstract

**Motivation:** Rapid development in long read sequencing and scaffolding technologies is accelerating the production of reference-quality assemblies for large eukaryotic genomes. However, haplotype divergence in regions of high heterozygosity often results in assemblers creating two copies rather than one copy of a region, leading to breaks in contiguity and compromising downstream steps such as gene annotation. Several tools have been developed to resolve this problem. However, they either only focus on removing contained duplicate regions, also known as haplotigs, or fail to use all the relevant information and hence make errors.

**Results:** Here we present a novel tool “purge_dups” that uses sequence similarity and read depth to automatically identify and remove both haplotigs and heterozygous overlaps. In comparison with the current standard, purge_haplotigs, we demonstrate that purge_dups can reduce heterozygous duplication and increase assembly continuity while maintaining completeness of the primary assembly. Moreover, purge_dups is fully automatic and can be easy integrated into assembly pipelines.

**Availability:** The source code is written in C and is available at https://github.com/dfguan/purge_dups.

## 1 Introduction

The superior throughput and increasing throughput of long read sequencing technologies, such as from Pacific Biosciences (Pacbio) and Oxford Nanopore Technologies (ONT), is revolutionizing the sequencing of genomes for new species (Phillippy, 2017). Long read assemblers, such as Falcon (Chin et al., 2016), Canu (Koren et al., 2017), miniasm (Li, 2016), typically generate haplotype-fused paths of a diploid genome, with Falcon-unzip (Chin et al., 2016) further able to separate the initial assembly into primary contigs and haplotigs. However, when there is high heterozygosity as in many outbred species, for example most insects and marine animals, the allelic relationships between haplotypic regions can be hard to identify, causing not only haplotigs to be mislabeled as primary contigs, but also overlaps to be kept among the primary contigs. The majority of these retained overlaps are between homologous chromosomes, and the resulting duplication harms downstream processes such as scaffolding and gene annotation, leading to incorrect results.

Tools such as Purge_haplotigs (Roach et al., 2018), Haplomerger (Huang et al., 2012), and the Redundans pipeline (Pryszcz and Gabaldón, 2016) have been designed to resolve this problem. Purge_haplotigs makes use of both read depth and sequence similarity to identify haplotigs. However, it does not identify heterozygous overlaps, and requires users to specify read-depth cutoffs manually. Both Haplomerger and Redundans seek to identify both haplotigs and overlaps, but they ignore read depth and rely only on the alignment of contigs to each other, which is prone to cause repetitive sequences to be over-purged.

Here we describe a novel purging tool, purge_dups, to resolve the haplotigs and overlaps in a primary assembly, using both sequence similarity and read depth. Purge_dups is now being used routinely in the Vertebrate Genomes Project assembly pipeline.

## 2 Methods

Given a primary assembly and long read sequencing data, we apply the following steps to identify haplotigs and overlaps. A more detailed description of the methods is available in the Supplementary Note.

2.1. We use minimap2 (Li, 2016) to map long read sequencing data onto the assembly and collect read depth at each base position in the assembly. The software then uses the read depth histogram to select a cutoff to separate haploid from diploid coverage depths, allowing for various scenarios where the total assembly is dominated by haploid or diploid sequence.
2.2. We segment the input draft assembly into contigs by cutting at blocks ‘N’s, and use minimap2 to generate an all by all self-alignment.
2.3. We next recognize and remove haplotigs in essentially the same way as purge_haplotigs, and remove all matches associated with haplotigs from the self-alignment set.
2.4. Finally we chain consistent matches in the remainder to find overlaps, then calculate the average coverage of the matching intervals for each overlap, and mark an unambiguous overlap as heterozygous when the average coverage on both contigs is less than the read depth cutoff found in step 1, removing the sequence corresponding to the matching interval in the shorter contig.

## 3 Results and Discussion

We describe the performance of purge_dups (v0.0.3) on three Falconunzip primary assemblies: *Arabidopsis thaliana* (At) (Chin et al., 2016), *Anopheles coluzzi* (Ac) (Kingan et al., 2019) and pinecone soldierfish *Myripristis murdjan* (Mm), and compare our results to those of purge_haplotigs (v1.0.4).

Purge_dups removes 96.4% of duplicated haploid-unique k-mers in the Falcon-Unzip assembly of Mm, compared to 81.2% removed by purge_haplotigs (Supplementary Figure 1). The corresponding numbers for At are 88.4% and 80.7% (Supplementary Figure 2). Supplementary Figures 3 and 4 show examples of regions where purge_dups removes both contained and overlapping duplication, whereas purge_haplotigs only removes fully contained duplication.

Table 1 presents results on assembly and completeness before and after purging for the three assemblies, using Benchmarking Universal Single-Copy Orthologs (BUSCOs) (Simão et al., 2015) to assess the consequences of purging for gene set completeness and duplication. The original draft primary assembly for At is longer than the TAIR10 reference (140Mb compared to 120Mb), and 6.2% of the genes are identified as duplicated. After being processed with purge_haplotigs, the assembly size decreases to 123 Mb, with 1.7% gene duplication. Purge_dups removes a further 2Mb and reduces the duplication rate to 1.1%, while keeping the same number of complete genes. For Ac the overall gene set completeness actually increases.

**Table 1:** BUSCO scores and assembly metrics

|  | BUSCO scores<sup>1</sup> (%) |  |  |  |  | Assembly size (Mb) | Num. Contigs | Ctg N50 (Mb) |
| --- | --- | --- | --- | --- | --- | --- | --- | --- |
|  | C | C(S) | C(D) | F | M |  |  |  |
| At-FU<sup>2</sup> | **98.1** | 91.9 | 6.2 | **0.3** | **1.6** | 140 | 172 | 7.96 |
| At-PD | 97.7 | **96.6** | **1.1** | 0.6 | 1.7 | 121 | 95 | 7.98 |
| At-PH | 97.7 | 96.0 | 1.7 | 0.6 | 1.7 | 123 | 109 | 7.98 |
| Ac-FU | 98.7 | 94.7 | 4.0 | 0.6 | 0.7 | 266 | 372 | 3.52 |
| Ac-PD | **98.8** | **98.5** | **0.3** | 0.6 | **0.6** | 246 | **190** | **4.03** |
| Ac-PH | **98.8** | 96.9 | 1.9 | **0.5** | 0.7 | 253 | 224 | **4.03** |
| Mm-FU | **95.8** | 79.0 | 16.8 | **2.0** | **2.2** | 1250 | 1290 | 2.63 |
| Mm-PD | 94.4 | 90.9 | 3.5 | 2.7 | 2.9 | 838 | 559 | 3.41 |
| Mm-PDS | 94.7 | **91.3** | **3.4** | 2.6 | 2.7 | 840 | **322** | **14.50** |
| Mm-PH | 94.5 | 89.1 | 5.4 | 2.4 | 3.1 | 888 | 517 | 3.29 |
| Mm-PHS | 94.7 | 87.5 | 7.2 | 2.5 | 2.8 | 891 | 481 | 3.48 |

1: C is for complete genes, C(S) for complete single copy genes, C(D) for complete duplicate genes, F for fragmented genes, and M for missing genes. 2: FU for Falcon-Unzip, PD for `Purge_dups`, PH for `Purge_haplotigs`, and PDS and PHS for `purge_dups` (respectively `purge_haplotigs`) after scaffolding and polishing.

For Mm, which has overall heterozygosity 1.1%, the effect is more dramatic. Assembly size is reduced from 1250 Mb to 840 Mb, and the gene duplication rate from 16.8% to 3.4%. Furthermore, after scaffolding with 10X Genomics linked reads using Scaff10x (https://github.com/wtsihpag/Scaff10X), the purge_dups assembly generated 208 scaffolds with N50 23.82 Mb, and gap filling within these scaffolds during polishing with Arrow closed a substantial number of gaps, increasing contig N50 from 2.63 Mb initially to 14.50 Mb. The scaffold and contig improvements were more modest when purge_haplotigs was used: 221 scaffolds with N50 8.17 Mb, and final contig N50 3.48 Mb. This indicates that divergent heterozygous overlaps can be a significant barrier to scaffolding, and that it is important to remove them as well as removing contained haplotigs. To take proper advantage of this, we recommend applying purge_dups directly after initial assembly, prior to scaffolding.

In conclusion, purge_dups can significantly improve genome assemblies by removing overlaps and haplotigs caused by sequence divergence in heterozygous regions. This both removes false duplications in primary draft assemblies while retaining completeness and sequence integrity, and can improve scaffolding. It runs entirely autonomously without requiring user specification of cutoff thresholds, allowing it to be included in an automated assembly pipeline.

## Acknowledgements

We thank members of the Vertebrate Genomes Project assembly group for input and advice, including Arang Rhie, Zemin Ning, William Chow, Ying Yan, Adam Phillippy and Erich Jarvis. The Mm genome was sequenced at the Sanger Institute as part of the Vertebrate Genomes Project, and we thank members of the Sanger Institute DNA pipelines group for generating the sequence data and Byrappa Venkatesh for providing the sample.

## Purge_dups supplementary note

### 1 Supplementary Methods

#### 1.1 Read depth cutoffs calculation

Given a read depth histogram *H*, purge_dups calculates the read depth cutoffs with the following algorithm.

- Initially calculate differences *H*′ = *H*_*i*+1_ – *H_i_* of *H*, then smooth these using their 10 nearest neighbours to approximate the local derivative.
- Next, use the smoothed derivatives to find the turning points.
- Next we consider two cases: (1) ≥ 2 maxima are found, or (2) single maximum.
- In case (1) we first merge local maxima and minima (within 3 bins). If following this merging there remain two maxima with a minimum in between then we take the minimum *v* as the threshold between haploid and diploid, with interval (*N, v*] for haploid and (*v*, 3*v*] for diploid, where *N* is the noise cutoff, user-configurable with default value 5. Otherwise we take the highest remaining maximum and drop into case (2).
- For case (2) we decide whether this single peak at *p* represents haploid or diploid depth by comparing it to the mean read depth. If the peak occurs at below mean read depth we consider it to be haploid and set the intervals as (*N*, 1.5*p*] for haploid and (1.5*p*, 4.5*p*] for diploid. If the peak is above the mean read depth then we take (*N*, 0.75*p*] for haploid and (0.75*p*, 2.25*p*] for diploid.

#### 1.2 Haplotypic duplication identification

Given a matching set of all versus all self alignments from minimap2, and read depth cutoffs from the previous section, purge_dups uses the following steps to identify the haplotypic duplications in a draft primary assembly:

1. Contained haplotig identification: purge_dups uses essentially the same way as purge_haplotigs to detect the contained haplotigs. If more than 80% bases of a contig are above the high read depth cutoff or below the noise cutoff, it is binned into the potential junk bin. Otherwise if more than 80% bases are in the diploid depth interval it is labelled as a primary contig, otherwise it is considered further as a possible haplotig. Next for each possible haplotig, we consider its best alignment to another contig. If its alignment score is larger than *s* (default 70) and max match score larger than *m* (default 200), it is marked as a repeat; if the alignment score is larger than *s* and max match score no larger than *m*, it is marked as a haplotig. Otherwise it is left as a candidate primary contig.
2. Haplotypic overlap identification: after purging the junk and contained haplotigs, purge_dups chains the matches between remaining candidate primary contigs to find collinear matches with the following process (Supplementary Figure 5):
  i. Given all matches between contig *Q* and contig *T*, purge_dups builds a direct acyclic graph (DAG) with the matches as vertices. Each vertex *V_i_* in DAG is denoted as a tuple (*s, e, h, t, d, m*), where *s* and *e* are the start and end position on *Q, h* and *t* are the start and end position on *T, d* is the orientation and *m* is the number of matched bases.
  ii. all vertices are ordered by their start positions on *Q*. For a pair of (*V_i_,V_j_*), suppose without loss of generality that *V_i_* is a predecessor of *V_j_*, they are both aligned in the forward direction, and the overlap between *V_i_* and *V_j_* on *Q* is represented by 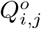, and on *T* is 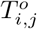. An edge exists between *V_j_* and *V_j_* if they meet the following conditions:

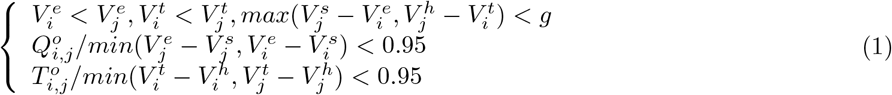 Where *g* is the maximum allowed gap size. Once the DAG is built, purge_dups will find the local optimal path by dynamic programming using the following recurrence equation:

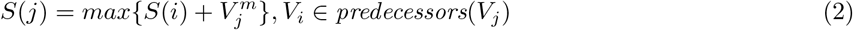

where *S_j_* is the score of *V_j_*.
  iii. After merging all the collinear matches, purge_dups filters out the nested matches and matches whose score is less than a threshold *l* (default: 10,000).
3. Calculate average read depth for the matching intervals in both the query and target, and only keep matches both of whose average read depths are below the diploid cutoff. Remove secondary and overlapping matches, defined as those for which the query region is contained within less than 85% of the matching region of another match from the same query, or no more than 85% of its sequence overlaps with another match. For remaining matches, move the sequence corresponding to the matching interval of the shorter contig into the haplotigs bin.

### 2 Supplementary Data

#### 2.1 Datasets

Datasets used in the experiments are listed as follows:

- At: We used the same assemblies for *Arabidopsis thaliana* as used in the purge_haplotigs paper, available at https://zenodo.org/record/1419699. SRA accessions for Pacbio reads are SRR3405291-SRR3405298 and SRR3405300-SRR3405326, and for paired-end Illumina reads are SRR3703081, SRR3703082, SRR3703105.
- Ac: The draft *Anopheles coluzzii* primary assembly that we used is available at https://drive.google.com/open?id=18osbKPOiUDWi65R5hpdbzNpGRgUtsJQy, the accession ID of the raw Pacbio reads is SRR8291675, and the RefSeq Accession ID of the AgamP4 PEST assembly for *Anopheles gambiae* is GCA_000005575.2.
- Mm: The draft Pacbio primary assembly is available at s3://genomeark/species/Myripristis_murdjan/fMyrMur1/assembly_cambridge/intermediates/falcon_unzip/fMyrMur1.PB.asm1.unzip.primary.fa.gz. All sequencing data are available in ENA with sample id SAMEA4872133.

#### 2.2 Software tools

The following software tools were used in the experiments:

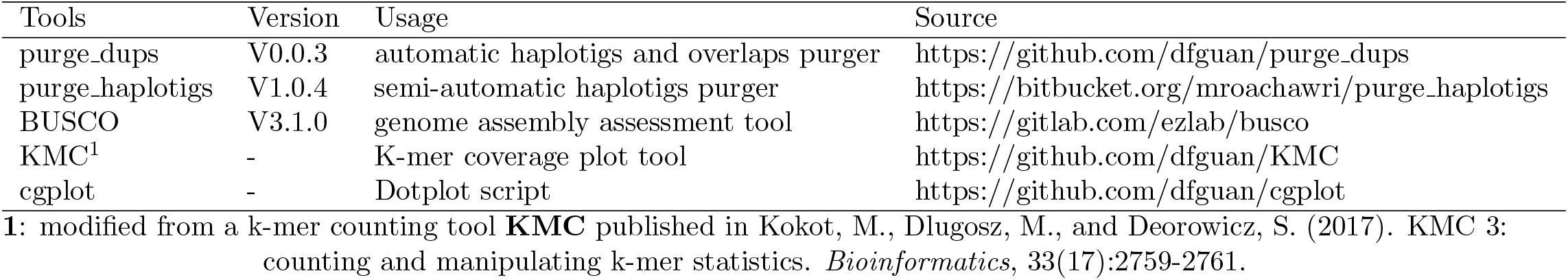

#### 2.3 Purge_dups commands

Given raw Pacbio reads alignment PAF files *pfs*, and a primary assembly *asm*. purge_dups use the following command to identify the haplotigs and overlaps:

~~~
   pbcstat $pfs // generates files PB.base.cov for base-level read depth and PB.stat for read depth histogram
   calcuts PB.stat > cutoffs 2>calcults.log
   split_fa $asm > $asm.split.fa
   minimap2 -xasm5 -DP $asm.split.fa $asm.split.fa > $asm.split.self.paf
   purge_dups -2 -T cutoffs -c PB.base.cov $asm.split.self.paf > dups.bed 2> purge_dups.log
  get_seqs dups.bed $asm > purged.fa 2> hap.fa
~~~

#### 2.4 Analysis parameters

Read depth cutoffs for purge_haplotigs were set manually and are shown here together with the databases used for BUSCO analysis:

| Assembly | Read depth cutoffs (low/middle/high) | BUSCO database |
| --- | --- | --- |
| At | 25/97/250 | embryophyta |
| Ac | 10/150/700 | diptera |
| Mm | 5/42/125 | actinopterygii |

### 3 Supplementary Figures

**Supplementary Figure 1:**
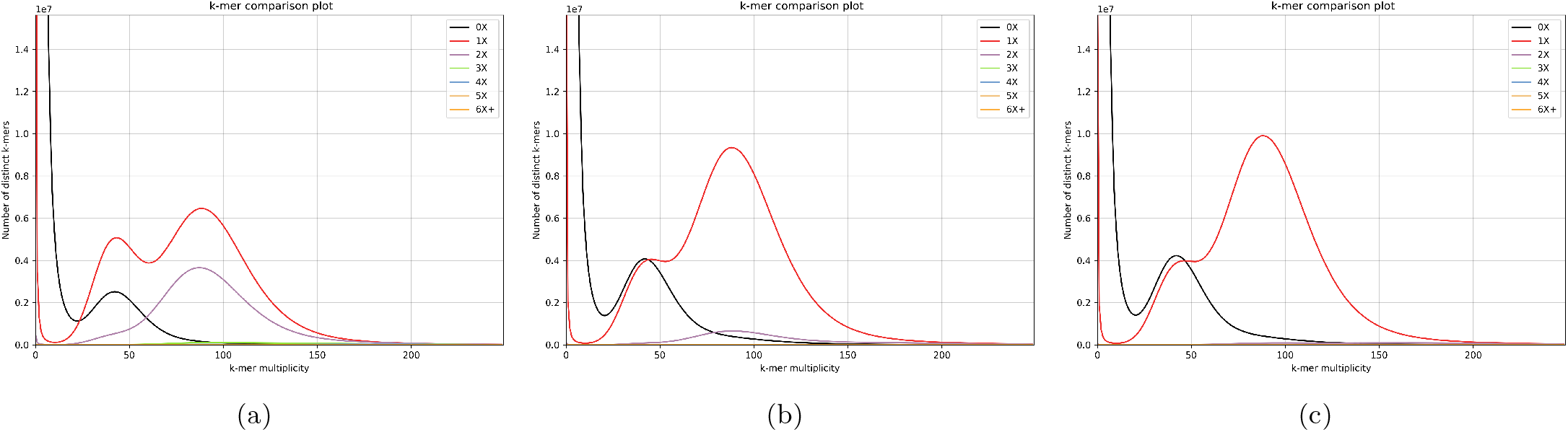
K-mer coverage plots for draft and purged Mm assemblies. The horizontal axis represents the copy number of k-mers in short reads from the same sample, the vertical axis shows the number of distinct k-mers, and the colored lines denote k-mers which occur the given number of times in the assembly. **(a)** The purple line shows 209.1 million 2-copy k-mers accumulating in the haploid and diploid areas, which correspond to duplicated haplotigs or overlaps in the primary assembly. **(b)** 39.3 million 2-copy k-mers remain after purging with purge_haplotigs. **(c)** Only 7.6 million 2-copy k-mers remain after purging with purge_dups.

**Supplementary Figure 2:**
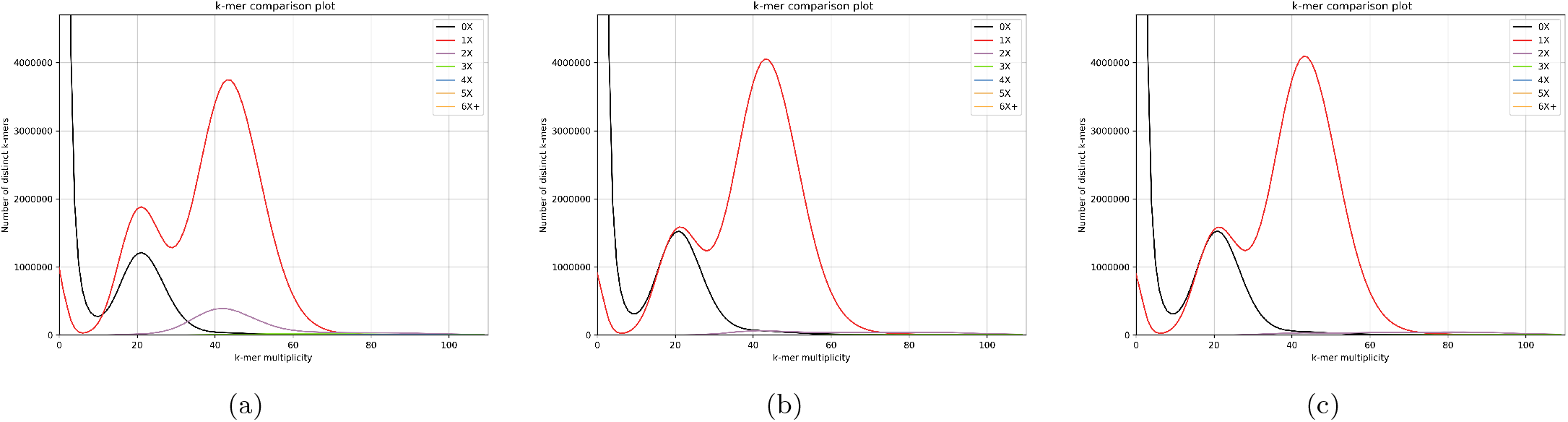
K-mer coverage plots for the At primary and purged assemblies. **(a)**: 8.06 million 2-copy k-mers remain in the diploid area of the original assembly (purple line). **(b)**: 1.56 million remain after purge_haplotigs. **(c)**: 0.94 million remain after purge_dups. We can not make this plot for assembly Ac because we do not have Illumina data from the same sample.

**Supplementary Figure 3:**
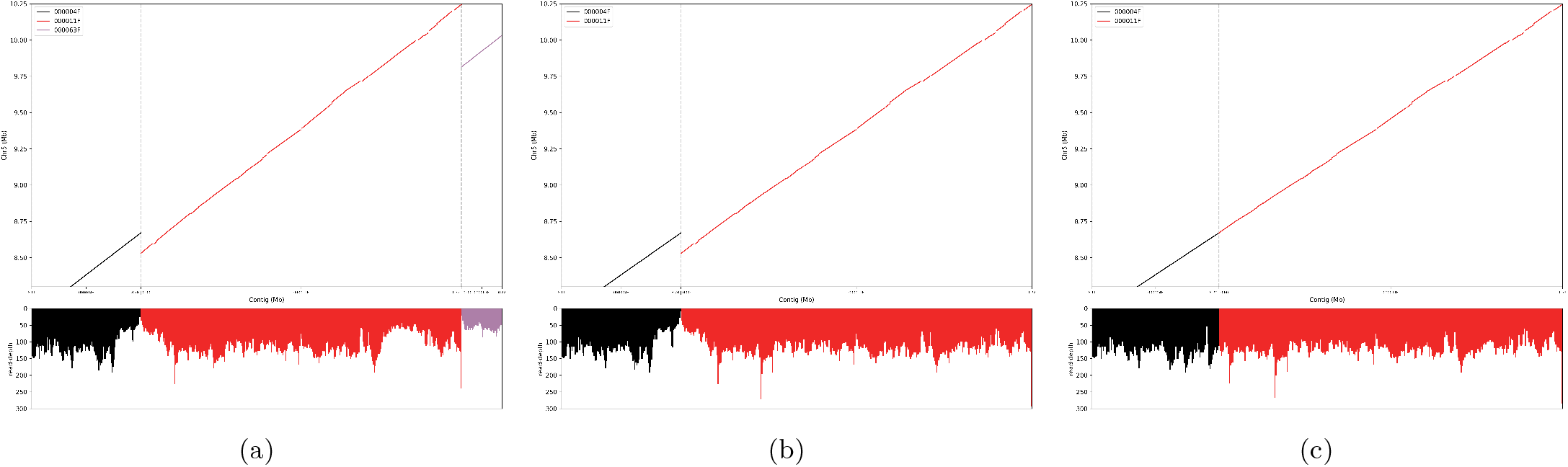
Dotplots of draft and purged At assemblies mapped to the TAIR10 reference genome. The horizontal axis represents the contigs in the assemblies, the upper vertical axis represents the reference chromosome, and the lower one shows the read depth for the contigs. **(a)** In the draft assembly, the right end of contig “000004F” and all of contigs “000011F” and “000063F” are aligned to part of chromosome 5. Contig “000063F” is contained in “000011F” and an overlap occurs at the ends of “000011F” and “000004F”. The read depth at the haplotypic and overlapped region drops to almost half of the diploid read depth (150). **(b)** In the purge_haplotigs assembly, the haplotig is removed, and read depth at the haplotypic region goes back to diploid read depth. However the overlap remains. **(c)** In the purge_dups assembly, both the haplotig and the overlap are removed and read depth goes back to normal across the whole range.

**Supplementary Figure 4:**
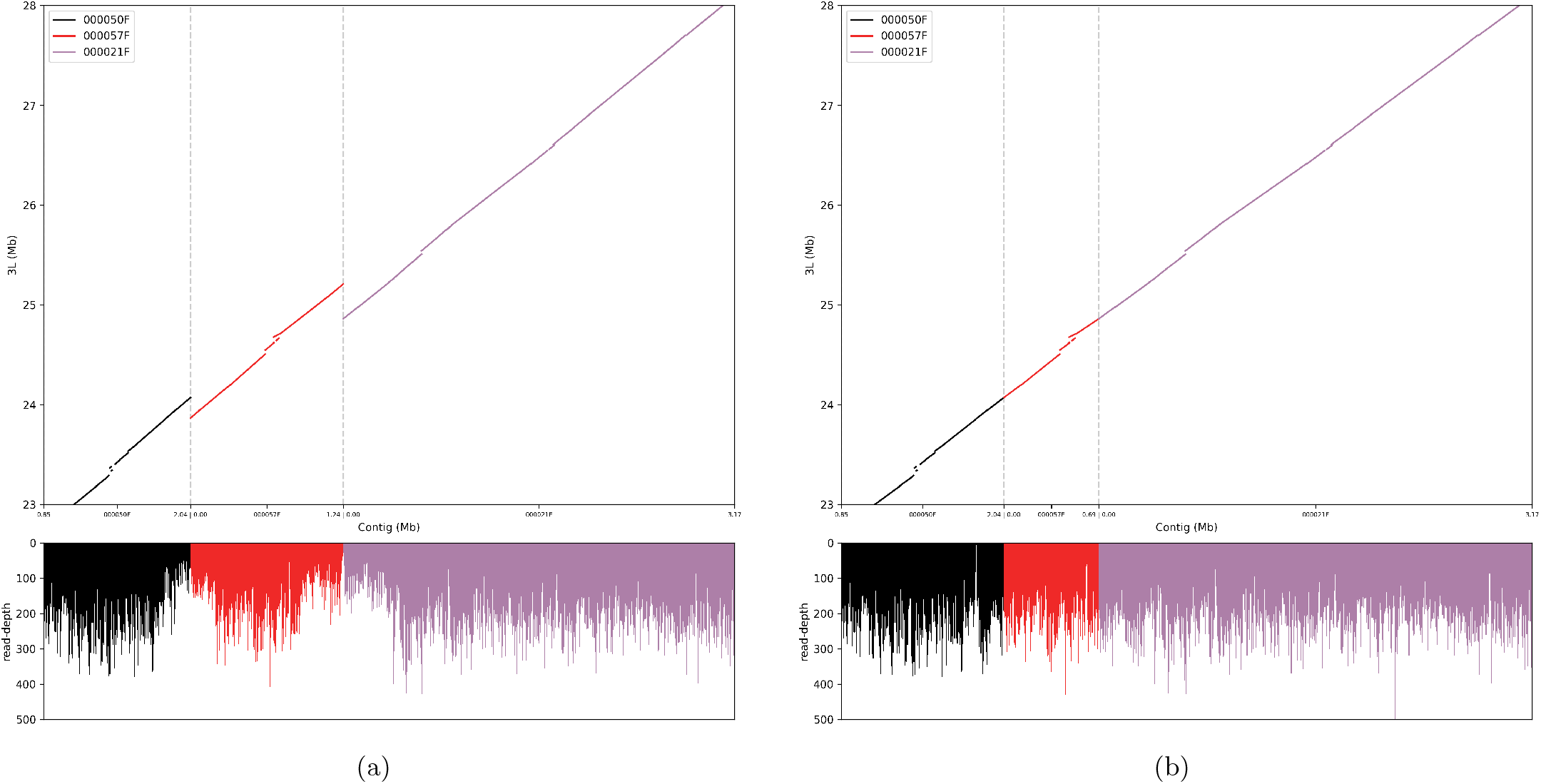
Dotplots on Ac draft primary and purge_dups assemblies. The horizontal axis represents the contigs in the assemblies, the upper vertical axis represents the reference chromosome, and the bottom one shows the read depth for the contigs. The draft and purged primary assemblies are mapped to AgamP4 PEST reference assembly. **(a)**: Contig 50F, 21F and 57F are aligned to 23-28 Mb region of chromsome 3L on PEST genome. Two overlaps are found, the read depth of the corresponding regions also drops to half of the normal diploid coverage. **(b)**: After purging with purge_dups, the overlaps are removed perfectly, and the read depth becomes even at the diploid level.

**Supplementary Figure 5:**
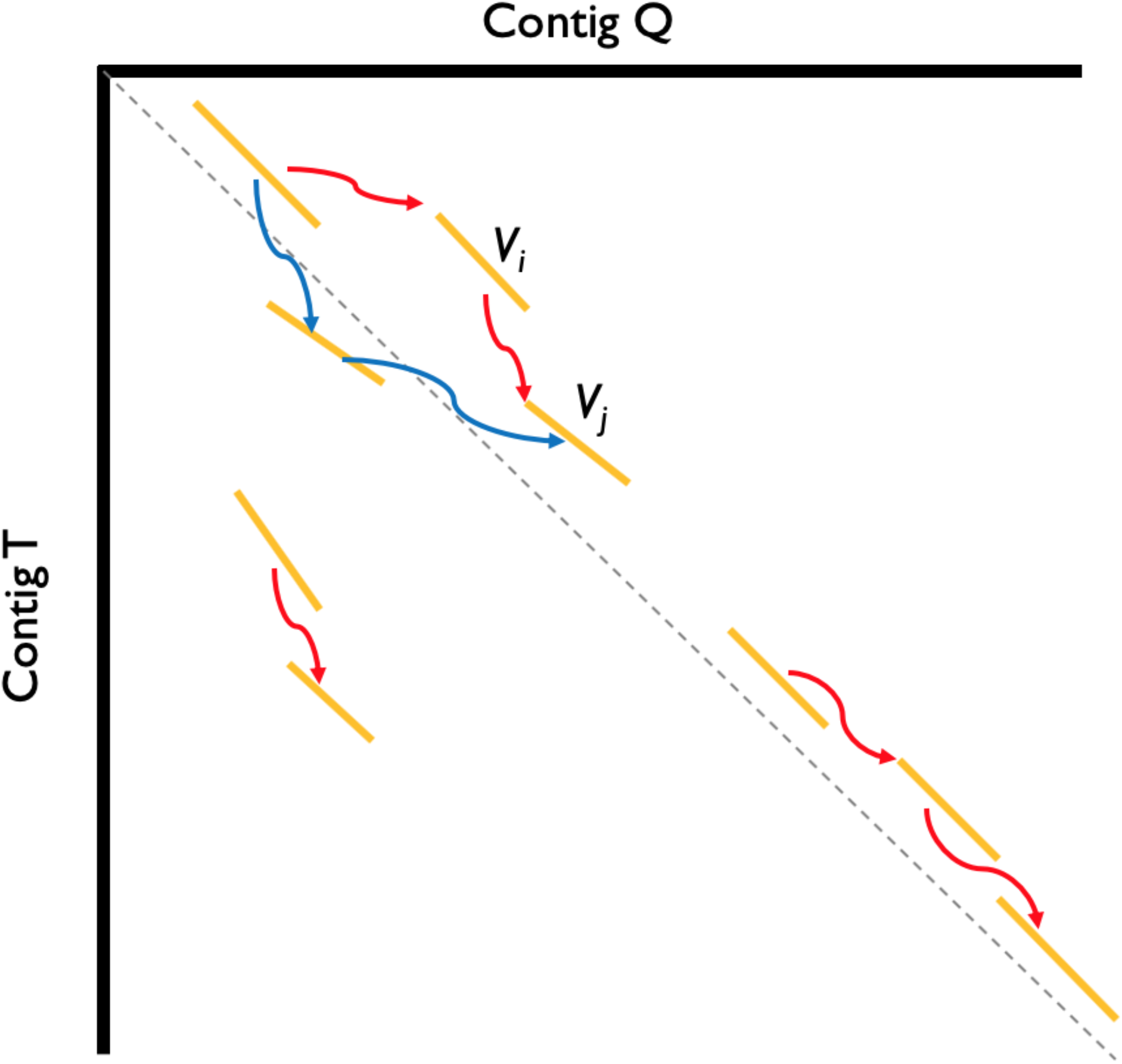
Illustration of chaining algorithm. Vertices representing the matches are the lines in orange, edges are shown in red and blue. Red edges are used to form a collinear match. Three collinear matching groups are found in this example.

